# E-cadherin regulates the stability and transcriptional activity of β-catenin in embryonic stem cells

**DOI:** 10.1101/2021.07.22.453344

**Authors:** Sinjini Bhattacharyya, Ridim D. Mote, Jacob W. Freimer, Surya Bansi Singh, Sandhya Arumugam, Yadavalli V. Narayana, Raghav Rajan, Deepa Subramanyam

**Affiliations:** National Centre for Cell Science, Ganeshkhind Road, Pune – 411007; SP Pune University, Ganeshkhind Road, Pune – 411007; Department of Genetics, Stanford University, Stanford, CA 94305, USA; Gladstone-UCSF Institute of Genomic Immunology, San Francisco, CA 94158, USA; Department of Microbiology and Immunology, University of California, San Francisco, San Francisco, CA 94143, USA; Indian Institute of Science Education and Research, Dr. Homi Bhabha Road, Pune -411008

**Keywords:** embryonic stem cells, E-cadherin, β-catenin, cell adhesion

## Abstract

E-CADHERIN is abundantly expressed in embryonic stem cells (ESCs) and plays an important role in the maintenance of cell-cell adhesions. However, the exact function of this molecule beyond cell adhesion, in the context of cell fate decisions is largely unknown. Using mouse ESCs (mESCs), we demonstrate that E-CADHERIN and β- CATENIN interact at the membrane and continue to do so upon internalization within the cell. Knockout of the gene encoding E-CADHERIN, *Cdh1*, in mESCs resulted in a failure to form tight colonies, accompanied by altered expression of differentiation markers, and retention of pluripotency factor expression during differentiation. Interestingly, *Cdh1^-/-^* mESCs showed a dramatic reduction in β-CATENIN levels. Transcriptional profiling of *Cdh1*^-/-^ mESCs displayed a significant alteration in the expression of a subset of β-CATENIN targets, in a cell-state dependent manner. While treatment with a pharmacological inhibitor against GSK3β could rescue levels of β-CATENIN in *Cdh1^-/-^* mESCs, expression of downstream targets were altered in a context-dependent manner, indicating an additional layer of regulation within this subset. Together, our results reveal the existence of a cell-state-dependent regulation of β-CATENIN and its transcriptional targets in an E-CADHERIN dependent manner. Our findings hint at hitherto unknown roles played by E-CADHERIN in regulating the activity of β-CATENIN in ESCs.

**Significance Statement:** Are cell adhesions only responsible for maintaining tissue architecture, or do they also regulate cell fate decisions during early embryonic stages by modulating the output of specific signalling pathways? In this study, we study the role of E-CADHERIN, a crucial component of cell-cell adhesions in the context of mouse embryonic stem cells (mESCs). We find that E-CADHERIN regulates the stability and activity of β-CATENIN in mESCs through physical interactions. However, the loss of E-CADHERIN affected the expression of only a subset of downstream targets of β-CATENIN in a cell-state dependent manner. This study highlights a critical cross-talk between molecules involved in cell-cell adhesion and the underlying signalling network critical for establishing cell fate during early mammalian development.

## Introduction

Cell-cell adhesion is required for the generation and maintenance of well-ordered three-dimensional tissues in higher organisms. Early events in embryonic development involve large-scale movements of sheets of cells, and tissue integrity is maintained by virtue of the well-coordinated action of cell-cell adhesion molecules (1). Among these, the best-studied are the classical cadherins, such as E-cadherin, which were first identified as trans-membrane proteins facilitating calcium-mediated cell-cell adhesion through the N-terminal extracellular domain (2, 3).

The importance of cadherins in maintaining tissue integrity was first demonstrated through experiments involving embryos expressing mutant forms of E-cadherin. These embryos displayed multiple phenotypes including dissociation of blastomeres and defects in epithelial integrity and gastrulation (1, 4, 5, 6). In the case of mouse embryos, E-cadherin facilitated compaction. However, E-cadherin null embryos could progress up to the implantation stage, presumably due to the maternal pool of E-cadherin (6). Studies from *Drosophila* and *C.elegans* also further demonstrated the crucial role played by cadherins in maintaining tissue integrity and cell-adhesion during morphogenetic movements – a process involving extensive forces (7, 8).

In addition to its role in cell adhesion, E-cadherin is known to interact through its C-terminal cytoplasmic domain with a range of proteins, notably the catenins: β- catenin, α-catenin and p120-catenin. β-catenin (CTNNB1), a key signal transducer in the Wnt pathway, exerts its action through translocation from the cytosol to the nucleus, where it binds to the TCF/LEF family of transcription factors to drive the expression of target genes (9–12). In the absence of active Wnt signalling, β-catenin undergoes phosphorylation by kinases including casein kinase I (CKI) and glycogen synthase-3β (GSK3β), marking it for proteasomal degradation (12–14) .

Cell culture based systems such as those involving embryonic stem cells (ESCs), isolated from the pre-implantation blastocyst, have emerged as popular model systems to study development and regeneration. These cells have the potential to differentiate and give rise to the cells belonging to all the three germ layers - ectoderm, mesoderm, and endoderm. E-CADHERIN is highly expressed in mouse ESCs (mESCs), and is essential for their ability to form compact colonies through intercellular junctions (15–17). Previous reports showed that *Cdh1^-/-^* mESCs displayed a scattered morphology, while retaining expression of pluripotency factors even in the absence of LIF (17). Subsequent studies however reported that reprogramming of somatic cells to a pluripotent state was complete only in the presence of *Cdh1*, with loss of *Cdh1* resulting in the differentiation of mESCs (15). Further, recycling of E-CADHERIN was found to be essential in the context of both human and mouse ESCs (16, 18). Recently it was also shown that β-catenin deficient mESCs could self-renew and retain expression of pluripotency markers (19). Interestingly, the expression of canonical Wnt targets was not altered in these cells, indicating that Wnt/β-catenin signalling may not be constitutively active in mESCs. In a parallel study, it was reported that β-catenin has repressive functions upon engagement with factors such as E2F6, HMGA2, and HP1ϒ, preventing the differentiation of mESCs towards a neural lineage. These findings reveal a new role for β-CATENIN whereby it not only activates the transcription of core pluripotency factors, but can also repress expression of genes associated with lineage differentiation to maintain the ground state of mESCs (20).

In spite of the extensive studies carried out on the individual roles of E-cadherin and β-catenin in context of mESCs, the significance of their interaction, inter- dependence, and effects on downstream targets has not been explored widely. In other words, are cell adhesion molecules just the structural glue, or do they also directly or indirectly regulate cellular outcomes and fate decisions, by modulating signalling pathway outputs? This manuscript attempts to address this question.

Here we demonstrate that E-CADHERIN interacts with β-CATENIN in mESCs even upon internalization. Constitutive loss of E-CADHERIN through the generation of *Cdh1^-/-^* mESCs result in their inability to form intact colonies and display a scattered morphology. Interestingly *Cdh1^-/-^* mESCs also display dramatically reduced levels of β-CATENIN and its associated activity. Treatment with a pharmacological inhibitor of GSK3β was able to rescue both the levels and activity of β-CATENIN. Interestingly, loss of E-CADHERIN resulted in the altered expression of a subset of β-CATENIN targets. This was further dependent on the differentiation state of the cells. While the expression of a subset of β-CATENIN targets could be rescued through GSK3β inhibition, this was heavily dependent on the status of *Cdh1* or the state of differentiation of the cell. Together, our results demonstrate that E-CADHERIN provides an additional layer for the regulation of β-CATENIN and its downstream targets in a cell-state dependent manner.

## Results

### E-CADHERIN and β-CATENIN continue to interact even upon internalization in mESCs

WT mESCs form compact, dome-shaped colonies by virtue of strong cell-cell adhesions (Supp. Fig. 1A). These cell-cell adhesions are largely facilitated by the cell adhesion molecule, E-CADHERIN (1). E-CADHERIN is known to bind β-CATENIN through a domain at its C-terminus, with β-CATENIN also playing a role in stabilising E-CADHERIN on the membrane (21). We found that a majority of E-CADHERIN and β-CATENIN predominantly co-localize at the cell membrane in mESCs (Fig. 1A). Cell adhesions are formed through the calcium-binding domains present on the extracellular region of E-CADHERIN. Treatment of mESCs with EGTA chelated Ca^2+^ causing disruption of cadherin-cadherin interactions and loss of cell-cell adhesions (Supp. Fig. 1A). This was accompanied by a translocation of both E-CADHERIN and β-CATENIN away from the membrane, into the cytoplasm (Fig. 1A), similar to what has been previously described in epithelial cells (22). Addition of excess Ca^2+^ re-established cadherin-cadherin interactions and cell-cell adhesion (Supp. Fig. 1A), along with a relocation of both E-CADHERIN and β-CATENIN back to the membrane (Fig. 1A). Interestingly, we found that β-CATENIN continued to co-localize and interact with E-CADHERIN when cell-cell adhesions were disrupted, and upon restoration of cell-cell adhesion with CaCl_2_ addition (Fig. 1A, 1B).

**Figure 1:**
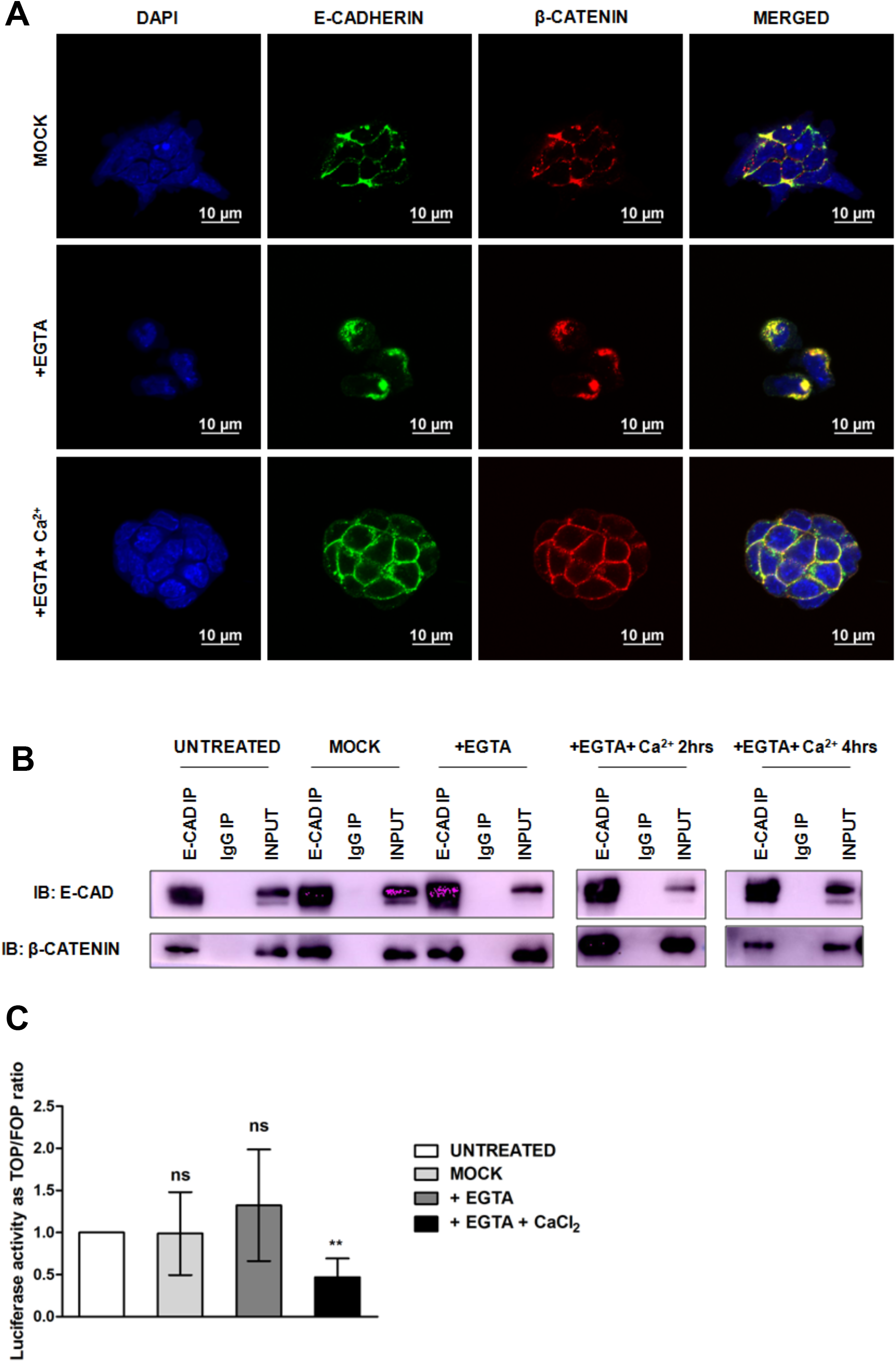
**E-CADHERIN continues to interact with β-CATENIN upon internalization in mESCs:** A. Representative confocal images showing localisation of E-CADHERIN and β- CATENIN in WT mESCs after treatment with EGTA or Ca^++^. Scale bar, 10µm. B. E- CADHERIN was immunoprecipitated from mESC extracts and blotted with anti-β-CATENIN monoclonal antibody after EGTA-induced disruption of E-cadherin mediated cell-cell adhesion and subsequent restoration by supplying Ca^++^. C. TOP Flash reporter assay showing level of β-CATENIN/TCF mediated transactivation in EGTA or Ca^++^ treated WT mESCs. Error bars represent mean±SD from 3 independent experiments; *p<0.05, **p<0.01, ***p<0.001 by two-tailed Student’s t- test.

In response to activation of the upstream Wnt pathway, β-CATENIN translocates to the nucleus, resulting in the transcriptional activation of downstream targets (22–24). Measurement of the transcriptional activity of β-CATENIN by the TOP-FLASH-based promoter-reporter assay showed that nuclear activity of the β-CATENIN/TCF complex was retained, and only minimally altered upon disruption of cell-cell adhesions in mESCs (Fig. 1C). Together this indicates that dislocation of E-CADHERIN from the cell membrane does not result in either destabilisation, or alteration of transcriptional activity of β-CATENIN, and that these two proteins continue to interact and remain as a complex even when forcefully internalized within the cell.

### *Cdh1^-/-^* mESCs display altered expression of pluripotency markers post differentiation

As E-CADHERIN and β-CATENIN continued to exist as a complex irrespective of the adhesion status of the cell, we further investigated the effect of *Cdh1* loss upon β- CATENIN levels in mESCs. Stable E-CADHERIN knockout (*Cdh1^-/-^*) mESCs were generated using CRISPR-Cas9 genome editing (Supp. Fig. 1B). *Cdh1^-/-^* mESCs were unable to form dome-shaped colonies, with cells presenting a scattered appearance (Supp. Fig. 1C), similar to what has been reported earlier (16, 18). *Cdh1^-/-^* mESCs had no detectable expression of E-CADHERIN (Supp. Fig. 1D), and had dramatically reduced *Cdh1* levels (Supp. Fig. 1E). We analysed the expression profile of *Cdh1^-/-^* mESCs compared to WT mESCs using RNA sequencing (RNA-seq). 2289 genes out of 15169 expressed genes were found to be differentially expressed (FDR adjusted p-value < 0.05). The volcano plot represents the genes whose expression was significantly altered as red dots (Fig. 2A). The expression of some of the highly upregulated and downregulated genes (indicated in Fig. 2A) was further validated using quantitative RT-PCR (Supp. Fig. 2A).

**Figure 2:**
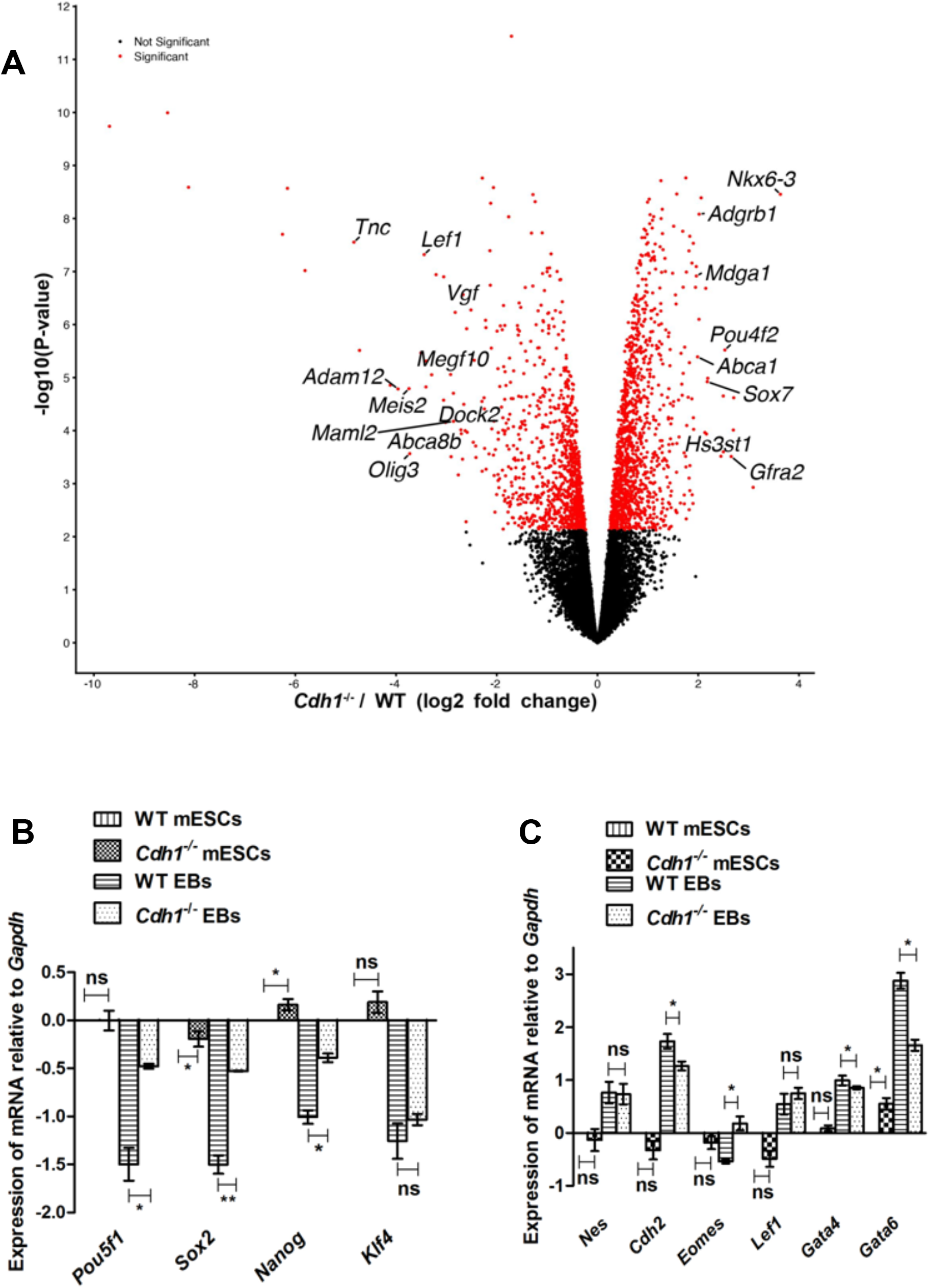
***Cdh1^-/-^* mESCs fail to downregulate the expression of pluripotency markers during differentiation**: A. Volcano plot depicting the genes exhibiting significant fold change in expression upon knocking out *Cdh1* in mESCs. B. Graph showing the expression of pluripotency markers in WT mESCs, *Cdh1^-/-^* mESCs, WT EBs and *Cdh1^-/-^* EBs. Expression is normalized to *Gapdh* and further normalized to levels seen in WT mESCs. C. Graph representing the expression of markers associated with the three germ layers (ectoderm, endoderm and mesoderm) at the transcript level in WT mESCs, *Cdh1^-/-^* mESCs, WT EBs and *Cdh1^-/-^* EBs. Expression is normalized to *Gapdh* and further normalized to levels seen in WT mESCs. For all experiments, error bars represent mean±SD from 3 independent experiments; *p<0.05, **p<0.01, ***p<0.001 by two- tailed Student’s t-test.

The RNA-seq data further indicated that the expression of most pluripotency markers were not significantly altered in *Cdh1^-/-^* mESCs compared to WT mESCs (Supp. Fig. 2B). Quantitative RT-PCR analysis of gene expression and immunocytochemistry further validated this observation (Fig. 2B, Supp. Fig. 2D). However, the expression of endoderm-specific genes such as *Gata4*, *Gata6*, *Nodal*, *Sox17* and *Sox7* were upregulated in *Cdh1^-/-^* mESCs as determined by RNAseq, with *Gata6* showing a significant increase in expression upon qRT-PCR validation (Supp. Fig. 2C, Fig. 2C). We further checked the expression of differentiation markers when embryoid bodies (EBs) were generated from *Cdh1^-/-^* mESCs. *Cdh1^-/-^* EBs retained the expression of *Pou5f1*, *Sox2* and *Nanog*, unlike their WT counterparts (Fig. 2B). However, they were still able to upregulate differentiation markers to an almost similar extent as WT EBs, with the exception of *Gata6*, indicating that their differentiation potential, while altered, was not entirely compromised (Fig. 2C). Therefore, we speculate that while the absence of E-CADHERIN does not completely alter the pluripotency profile of mESCs, it affects the downregulation of pluripotency marker expression during differentiation.

### E-CADHERIN regulates the stability of β-CATENIN in mESCs

β-CATENIN has long been known to interact with E-CADHERIN and play a role in stabilising it at the membrane (21). Previously, we have shown that both proteins exist in a complex irrespective of the adhesion status of the cell (Fig. 1). *Cdh1^-/-^* mESCs showed a significant decrease in β-CATENIN protein levels (Fig. 3A, Supp. Fig. 3A), although there was no significant change at the transcript level (Fig. 3B). This was also accompanied by a significant decrease in β-CATENIN-dependent transcriptional activity in *Cdh1^-/-^* mESCs (Fig. 3C). To determine whether the decrease in β-CATENIN was dependent entirely upon the loss of E-CADHERIN we overexpressed either the full-length WT E-cadherin (*Cdh1*FL), or a mutant E-cadherin lacking the C-terminal, β-CATENIN binding site (*Cdh1Δβctn*) (Fig. 3D). Rescue of β-CATENIN expression was observed only in the presence of *Cdh1*FL, and not upon overexpression of the truncated version, *Cdh1Δβctn* (Fig. 3E, Supp. Fig. 3B). Previous studies suggest that the interaction of the cadherin cytoplasmic tail with catenins mediate the formation of cell-cell adhesion in the embryonic state (4). Moreover, it was also established that this catenin-cadherin interaction depends on the phosphorylation status of their interacting domains (25, 26). From our results it is clear that the reverse is also true where E-CADHERIN plays a major role in stabilising β-CATENIN levels, and its activity, presumably through interactions with its cytoplasmic domain in mESCs.

**Figure 3:**
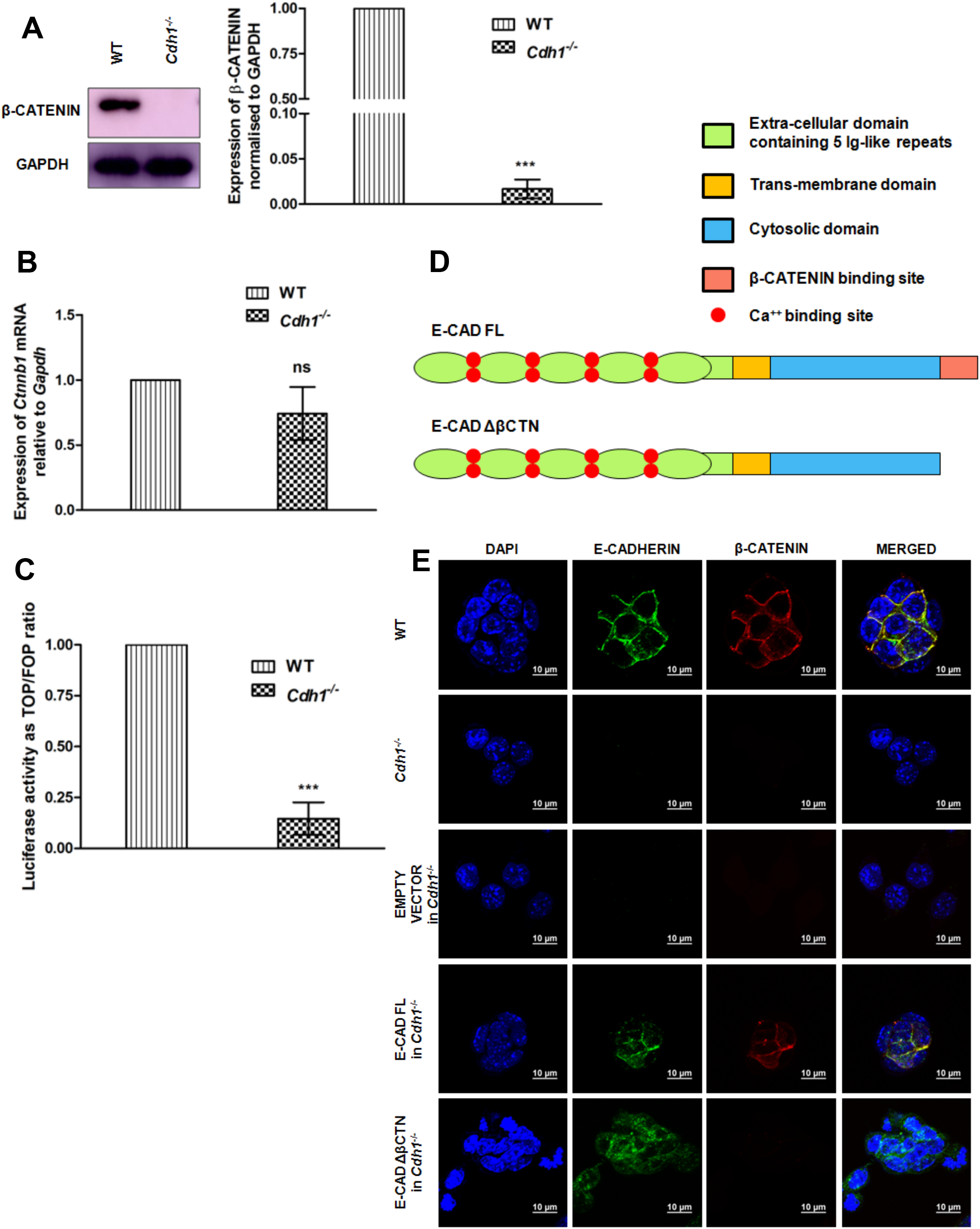
**Interaction with E-CADHERIN regulates the stability of β-CATENIN in mESCs:** A. Western blot showing level of β-CATENIN expression in *Cdh1^-/-^* mESCs compared to WT; graphical representation for the densitometric analysis of the blot is provided to the right. B. Graph showing levels of *Ctnnb1* in *Cdh1^-/-^* mESCs normalized to *Gapdh*. C. TOP-Flash reporter assay showing levels of β- CATENIN/TCF mediated transactivation in *Cdh1^-/-^* mESCs compared to WT. D. Schematic representation of WT E-cadherin (E-CAD FL) and the mutant form which lacks the β-CATENIN binding site (E-CAD ΔβCTN). E. Immunofluorescence images of stably transfected *Cdh1^-/-^* mESCs expressing E-CAD FL or E-CAD ΔβCTN and endogenous β-CATENIN. Untransfected WT mESCs, untransfected *Cdh1^-/-^* mESCs and *Cdh1^-/-^* mESCs stably transfected with empty vector were used as controls. Scale bar, 10µm. For all experiments, error bars represents mean±SD from 3 independent experiments; *p<0.05, **p<0.01, ***p<0.001 by two-tailed Student’s t- test.

### Pharmacological inhibition of GSK3β can rescue β-CATENIN levels and activity in *Cdh1^-/-^* mESCs

In addition to existing as a part of the cell adhesion complex at the plasma membrane where it interacts and stabilises E-CADHERIN, β-CATENIN also plays an important role in the Wnt signalling pathway driving the expression of numerous target genes involved in regulating multiple cellular processes. When the Wnt signalling pathway is inactive, β-CATENIN is phosphorylated by GSK3β marking it for ubiquitination and subsequent proteasomal degradation. We speculated that the loss of β-CATENIN observed upon depletion of E-CADHERIN may be due to its destabilisation in a GSK3β-dependent manner. Therefore, we investigated if β-CATENIN levels in *Cdh1^-/-^* mESCs could be rescued by inhibiting GSK3β-mediated phosphorylation using a pharmacological inhibitor, CHIR99021. We observed a dose-dependent rescue of β-CATENIN in *Cdh1^-/-^* mESCs upon treatment with CHIR99021 (Fig. 4A), accompanied by an increase in its transcriptional activity measured using the TOP-Flash reporter assay (Fig. 4B). Together, these data indicate that depletion of E-CADHERIN targets β-CATENIN for phosphorylation by GSK3β leading to its destabilization in mESCs. To determine whether similar mechanisms also operated during the process of differentiation, we determined the levels of E-CADHERIN and β-CATENIN in embryoid bodies (EBs). WT EBs exhibited a downregulation in E-CADHERIN levels compared to mESCs (Supp. Fig. 3C). Interestingly, β-CATENIN was upregulated in the *Cdh1^-/-^* EBs compared to *Cdh1^-/-^* mESCs which lacked detectable β-CATENIN (Supp. Fig. 3C), presumably through an upregulation of *Cdh2* (Fig. 2C), which may function to compensate for the loss of *Cdh1.* Our data therefore suggests that while β-CATENIN levels and stability may be entirely dependent on E-CADHERIN in mESCs, alternate mechanisms, independent of E-CADHERIN, may exist during differentiation.

**Figure 4:**
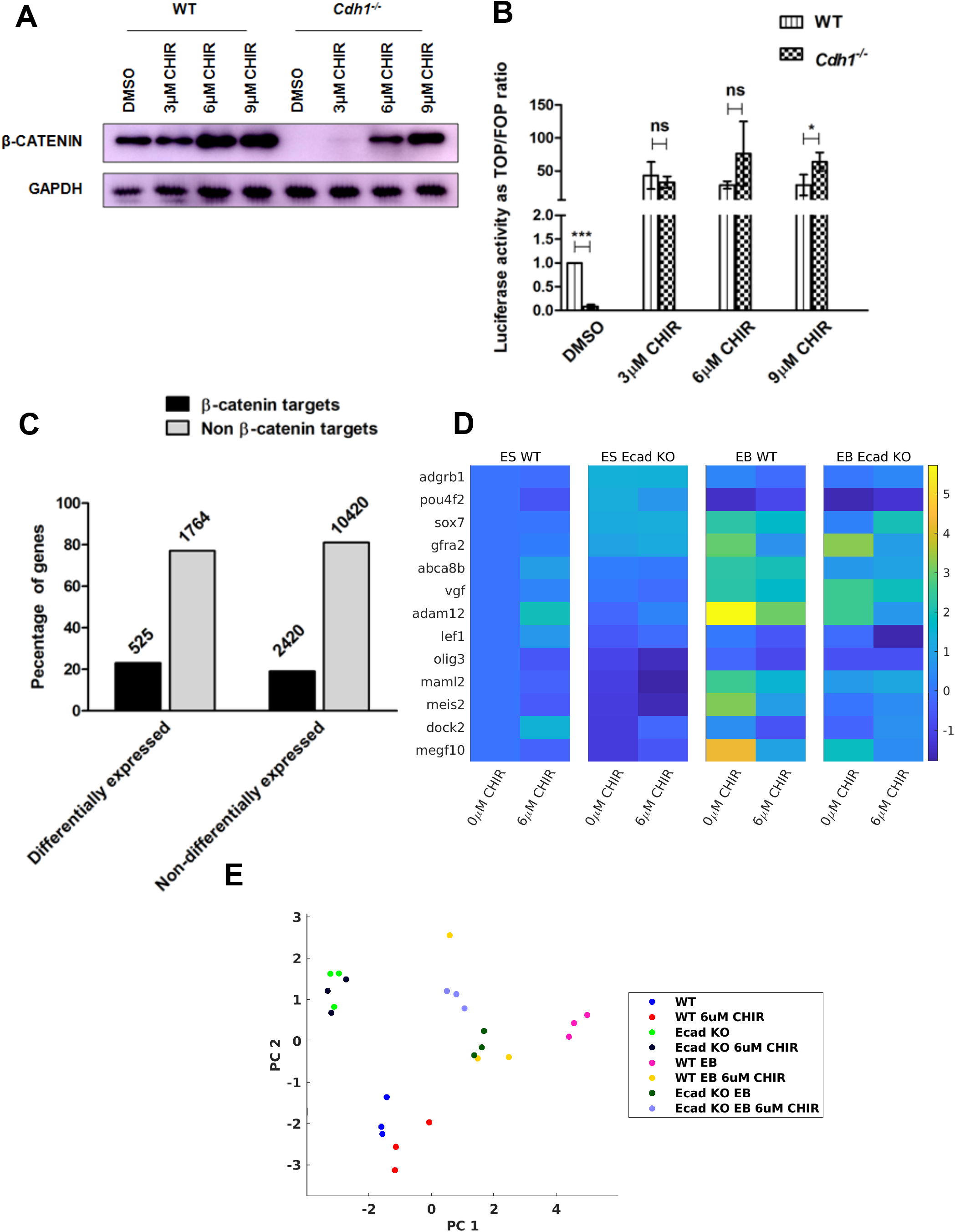
**E-CADHERIN and β-CATENIN are involved in complex interactions to regulate downstream targets:** A. Western blot showing levels of β-CATENIN in WT and *Cdh1^-/-^* mESCs after treatment with GSK3β inhibitor, CHIR99021 (CHIR) at the indicated concentrations. DMSO was used as control. B. TOP Flash reporter assay showing levels of β- CATENIN/TCF mediated transactivation in *Cdh1^-/-^* mESCs compared to the WT following treatment with CHIR at the indicated concentrations. Error bars represent mean±SD from 3 independent experiments; *p<0.05, **p<0.01, ***p<0.001 by two- tailed Student’s t test. C. Graph showing that 525 (∼23%) of the differentially expressed genes and 2420 (∼19%) of the non-differentially expressed genes from the RNA-seq analysis were β-CATENIN targets. D. Heat map showing the expression of 13 targets of β-CATENIN. Expression levels are relative to *Gapdh* and normalized to wild type. Colours represent the log of expression levels. E. Plot showing log-transformed gene expression levels for 13 targets of β-CATENIN projected onto the first two principal component axis. Individual points represent individual experiments (n=3). Colours represent different experimental conditions.

### Regulation of expression of β-CATENIN targets in *Cdh1^-/-^* mESCs

The significant reduction in the activity of β-CATENIN/TCF in *Cdh1^-/-^* mESCs prompted us to check the expression of β-CATENIN targets in these cells. We overlaid the ChIP-seq data published by Tao *et al* (20) with our RNA-seq data to analyse the expression profile of *bona fide* β-CATENIN targets. 23% of differentially expressed genes (both up- and down-regulated) were β-CATENIN targets (Fig. 4C), indicating that β-CATENIN may not only act as an activator, but also as a repressor in mESCs. 19% of the non-differentially expressed genes were also β-CATENIN targets (Fig. 4C), suggesting that there may be additional mechanisms working to maintain their expression.

We next asked whether the expression of differentially regulated β-CATENIN targets were influenced by the loss of *Cdh1* and/or the state of differentiation and whether changes in expression, if any, under both these conditions could be rescued by inhibiting GSK3β. We hypothesized that since *Cdh1^-/-^* mESCs have reduced levels of β-CATENIN, the expression of its targets should also be reduced. Additionally, based on our observation that GSK3β inhibition could restore β- CATENIN levels, we hypothesized that this should also be able to restore β- CATENIN target expression. We examined the expression levels of a subset of β- CATENIN targets in WT mESCs, *Cdh1^-/-^* mESCs, WT EBs and *Cdh1^-/-^* EBs, each in two different conditions; with 6µM CHIR99021 or with vehicle control (DMSO/ 0µM CHIR99021) (Supp. Fig. 4). While the expression of a majority of genes were affected by the absence of *Cdh1* in mESCs and EBs, the application of CHIR99021 did not restore gene expression in the case of all target genes (Fig. 4D; compare KO 6µM CHIR99021 with WT; Fig. 4E, Supp. Fig. 4). This suggested the existence of a complex interplay involving the presence/absence of *Cdh1*, cell state and CHIR99021 treatment. To understand this better, expression levels of individual genes were log-transformed and fit using linear mixed effect models with *Cdh1* status (WT or KO), differentiation state (ES or EB) and inhibitor concentration (0 or 6µM) as predictors (Supp Table 1). In addition, we also considered all possible 2- way interactions between these predictors and a 3-way interaction between all 3 predictors. Expression of individual genes were very well fit by these models (median r^2^ – 0.88; range – 0.41 – 0.98). For 12/13 genes, interaction terms were strong predictors of gene expression, demonstrating context-specific changes in levels (Supp. Fig. 4, Supp. Table 1 and Fig. 4D). For eg. the effect of CHIR on *Gfra2* expression was differentiation-state dependent; 6µM of CHIR increased expression level in mESCs, but not in EBs, irrespective of *Cdh1* status. For 6/13 genes, the 3- way interaction was a signficant predictor of expression level. For eg. *Maml2* – 6µM CHIR reduced expression levels in ES cells independent of *Cdh1* status. However, in EBs, 6µM CHIR reduced expression levels of *Maml2* only in WT and not in *Cdh1^-/-^* cells.

To better understand general patterns of gene expression across different conditions, we used dimension reduction techniques. Specifically, for each gene (n=13 genes), we converted log expression levels into z-scores by normalizing across all experimental conditions. Each biological repeat from different experimental conditions was represented by a vector with normalized expression of the 13 genes. We then used principal components analysis for visualizing differences in gene expression across experimental conditions (Fig. 4E, Supp. Table 2). The first two principal components explained 67.97%, and the first six principal components explained 94.5% of the variance in the data (Supp. Table 2). Projection of the data onto the first two principal components also revealed context-specific differences in gene expression. *Cdh1* status and differentiation state were clear predictors of expression levels as seen by the clustering of data (Fig. 4E). Inhibiting GSK3B using 6µM CHIR was context-specific and depended on both *Cdh1* status and differentiation state of the cell. Specifically, CHIR effects were minimal in ES cells independent of *Cdh1* status. However, in EBs, CHIR effects depended on the *Cdh1* status; effects were observed when *Cdh1* was present and effects were minimal when *Cdh1* was knocked out. Biological repeats for each experimental condition clustered together in this space, validating the use of this analysis to understand general patterns of gene expression across conditions. In summary, our results indicate that there exist complex interactions between E-CADHERIN and β- CATENIN in the regulation of β-CATENIN targets which cannot simply be rescued just by stabilising it in the absence of E-CADHERIN through modulation of GSK3β, but may be influenced and dependent on cell state.

## Discussion

The importance of cadherin-mediated cell-cell adhesions have been demonstrated in a wide range of biological scenarios ranging from embryonic development to cancer and metastasis. However, the role of cadherins in regulating signalling in the context of stem cells remains relatively unexplored. Here we show that E-CADHERIN interacts with β-CATENIN to regulate its stability and signalling capacity in a cell state dependent manner. We demonstrate that similar to epithelial cells, β-CATENIN and E-CADHERIN interact at the membrane and continue to do so upon internalization. The complex containing E-CADHERIN and β-CATENIN has previously been shown to be important for differentiation of mESCs in the absence of LIF (7). We find that *Cdh1^-/-^* mESCs maintain expression of core pluripotency markers, while also expressing markers of the endodermal lineage. When subjected to differentiation, *Cdh1^-/-^* mESCs retained expression of pluripotency markers, indicating an indirect role played by E-CADHERIN in regulating cell fate. Interestingly, *Cdh1^-/-^* EBs expressed sufficient levels of β-CATENIN, presumably through the stabilizing action of other cadherins, such as N-CADHERIN (CDH2), which is upregulated during differentiation.

Loss of E-CADHERIN resulted in the destabilization of β-CATENIN in mESCs, in a GSK3β-dependent manner. Since the absence of E-CADHERIN caused a significant decrease in the transcriptional activity of β-CATENIN, we expected a decrease in the expression of its downstream targets. However, the expression of a number of target genes increased while many remained unchanged. This was in line with observations reported by Tao et al. where they showed that β-CATENIN can repress transcription of lineage-specific genes in a TCF3-independent manner (20). Interestingly, while β-CATENIN levels in *Cdh1^-/-^* mESCs were restored upon treatment with a GSK3β inhibitor, the expression of downstream targets was not rescued, indicating that the level of β-CATENIN did not directly correlate with the expression of downstream targets. Additionally, the expression profile of β-CATENIN target genes was completely different during the course of differentiation in *Cdh1^-/-^* EBs compared to WT EBs.

Our observations raise a number of possibilities. Firstly, E-CADHERIN may function as a hub to allow other proteins to interact and regulate the activity and stability of β- CATENIN. In the context of colon cancers, E-CADHERIN has been shown to bind a large number of proteins (28). A detailed identification of the proteins that E-CADHERIN binds to in ESCs and their interaction to regulate downstream signalling needs to be undertaken. Further, while other cadherins, such as N-CADHERIN (*Cdh2*) may be capable of stabilizing β-CATENIN, this may be insufficient to drive the expression of downstream targets in the context of *Cdh1^-/-^* mESCs.

Secondly, β-CATENIN can act as both an activator and repressor of downstream targets. A similar conclusion was also drawn based on the study from Tao et al. (20) and this may be dependent on the influence and presence of additional interactors. Thirdly, GSK3β may not influence the expression of all targets of β-CATENIN, indicating that additional signals may be required to drive downstream target transcription. Recent work using Xenopus embryos has demonstrated that Dkk2 can influence neural crest cell activity by driving Wnt/ β-catenin signalling in a GSK3β- independent manner (29). Finally, it is possible that the loss of E-CADHERIN may not result in the destabilization of all forms of β-CATENIN, and that a small, but undetectable fraction of β-CATENIN may linger, enough to drive the expression of a number of downstream targets that remain unchanged in *Cdh1^-/-^* mESCs. This begs the question of whether different pools of β-CATENIN exist, some of which are dependent on interactions with E-CADHERIN while others are not. Recent work coupling live cell microscopy to computational modeling describes the dynamics of various pools of β-CATENIN in HAP1 cells (30). However, it is unknown whether similar pools exist in mESCs and if these respond to loss of E-CADHERIN and /or treatment with inhibitors to GSK3β. Each of these possibilities may further be inter- dependent on the other, providing a complex network that drives and regulates expression of genes during fate determination.

While our studies have mostly focussed on the E-CADHERIN/ β-CATENIN complex, it must also be considered that E-CADHERIN interacts with p120 which contributes to its stability at the membrane (31). It is therefore a possibility that the loss of E-CADHERIN may also affect the spatial distribution of p120. Loss of p120 in mESCs is known to cause an increase in the expression of pluripotency markers such as *Oct4* and *Sox2* and a decrease in the expression of differentiation marker genes. Differentiation towards the endodermal lineage is also specifically hampered in *Ctnnb1^-/-^* EBs (32). Moreover, p120 is also reported to regulate the balance between pluripotency and differentiation by promoting degradation of the REST-CoREST complex, a transcriptional repressor of neural differentiation genes (33). Therefore, it would be interesting to interrogate the impact on the pool of p120 in *Cdh1^-/-^* mESCs and EBs.

Together, our study opens up new avenues for understanding the role of cell-cell adhesions in regulating downstream signalling pathways during cell fate determination in a context-dependent manner.

## Materials and methods

### Cell culture

WT V6.5 mESCs and *Cdh1^-/-^* V6.5 cells were cultured in the absence of feeders on 0.2% gelatin-coated plastic tissue culture dishes (Corning). Cells were replated at a density of 2×10^3^ cells/cm^2^ every 3 days after dissociation with 0.25% trypsin–EDTA (1X) (Gibco, cat no. 25200-056). Cells were grown in Knockout DMEM (Gibco, cat no.10829-018) containing 15% FBS (Gibco, cat no. 10270-106), 2mM L-glutamate 100X (Gibco, cat no. 25030-081), 1X Penicillin/streptomycin (Gibco, cat no. 15140-122), 1mM of 100X MEM non-essential amino acids (Gibco, cat no. 11140-050), 2-mercaptoethanol 1000X (Gibco, cat no. 21985-023) and 1000U LIF (Leukemia Inhibitory Factor) prepared in-house.

HEK293T cells were grown on plastic cell culture dishes (Corning) and replated at a density of 1×10^4^ cells/cm^2^ every 2 days after trypsinization. Cells were grown in DMEM (Gibco, cat no. 11960-044) containing 10% FBS, 1X Penicillin/streptomycin. Cells were grown in aseptic incubators at 37°C with 5% CO_2_.

Additional materials and methods are described in detail in Supplementary Information.

## Data sharing

Reagents will be available upon request to

## Author contributions

D.S and S.B conceived and designed the study. S.B. carried out experiments described in Figs. 1, 3A,C-E, 4A,B; Supp. Fig. 1A,C,D, 2D, 3. S.B and R.D.M. contributed to experiments in Fig. 2B,C, 3B, 4D,E; Supp. Fig. 1E, 2A. J.W.F. did the analysis for Fig. 2A, 4C; Supp. Fig. 2B,C. S.B.S helped with imaging for experiments described in Fig. 1A, Supp. Fig. 2D . S.A helped with experiments 1B and Supp. Fig. 3B,C. Y.V.N made the Cdh1^-/-^ mESCs as described in Supp. Fig. 1B. R.R. analysed data for Fig. 4D,E; Supp. Fig. 4; Supp. Table 1,2. D.S. and S.B. wrote the manuscript with help from R.R., R.D.M. and J.W.F. All authors reviewed and approved the manuscript.

## Acknowledgements

This work was supported by funds to D.S. from the Wellcome Trust – Department of Biotechnology (DBT) India Alliance (Intermediate Fellowship- IA/I/12/1/500507), and NCCS intramural funding. S.B. is a recipient of a Senior Research Fellowship from CSIR, India; S.B.S is a recipient of a Senior Research Fellowship from the Department of Biotechnology, India; JWF is funded by NIH grant R01HG008140; R.R is funded by Ramalingaswami fellowship from DBT, India (BT/HRD/35/02/2006). We thank members of the Subramanyam lab for constructive discussion.

## Supplementary Information

### Materials and Methods

#### CRISPR/Cas9 targeting

A single guide RNA (sgRNA) was designed to target the second exon of *Cdh1* gene and cloned into the pX459 vector (Addgene, cat no.62988). 2µg of the cloned plasmid was transfected into 5×10^3^ mESCs using Lipofectamine 2000 as per manufacturer’s instructions (Invitrogen, cat no.11668- 019). 24 hours post transfection, cells were put under antibiotic selection using 1µg of puromycin/ml of media for 48 hours. Surviving cells were replated on 0.2% gelatin-coated 96-well tissue culture plates (Corning) at single-cell density per well. After 5 days, the cells were screened based on their scattered morphology. Knockout clone (*Cdh1^-/-^*) was confirmed by Western blotting analysis.

#### Lentivirus production and infection

cDNAs encoding full length E-cadherin (E- CAD FL), and E-CADHERIN lacking the C-terminal site for β-CATENIN binding (E-CAD ΔβCAT) were amplified using specific primers (Supp. Table 3) and cloned into the EcoRI and NotI restriction sites of the lentiviral expression vector pCDH-EF1- FHC (Addgene, no. 64874). To generate lentiviruses, pCDH-EF1-FHC containing the specific E-CADHERIN variant, psPAX2 (Addgene, no. 12260), and pMD2.G (Addgene, no. 12259) were co-transfected into 60% confluent HEK293T cells using FuGENE HD transfection reagent (Promega, cat no. E2311). The viral supernatant was collected 48 hours post transfection and filtered through 0.45µm filter (Pall Life Sciences, cat no. 4654). 5×10^5^ *Cdh^-/-^* mESCs were infected with 1ml of viral supernatant in the presence of 5µg/ml Polybrene. Infected cells were subjected to antibiotic selection with 1µg puromycin/ml in media for 48 hours. Surviving cells were expanded and analysed by Western blot to check for expression of E-CADHERIN.

#### Chemical treatment

EGTA (Sigma, cat no. E8145) at pH 8.0 was used to internalise E-CADHERIN at a final concentration of 4mM, calcium chloride was used in fresh medium for the re-establishment of cell-cell adhesion mediated by dimerisation of E-CADHERIN at a final concentration of 1.8mM. GSK3β inhibitor CHIR99021 (SIGMA, cat no. SML1046) was used at concentrations of 3µM, 6µM or 9µM.

#### Antibodies

Primary antibodies used were against E-CADHERIN (BD Bioscience, cat no.610182, mouse), non-phospho active β-CATENIN (Ser33/37/Thr41) (D13A1) (CST, cat no. 8814, rabbit), and GAPDH (14C10) (CST, cat no. 2118, rabbit) at specific concentrations.

#### Co-immunoprecipitation

Cells were lysed in 1% NP40 lysis buffer. Co-immunoprecipitation was performed by incubating cell lysate with Dynabeads ProteinA (Invitrogen, cat no. 10001D) coated with specific antibody overnight. Normal mouse (G3A1) mAb IgG1 isotype control (CST, cat no. 5415) was used as antibody control. After incubation, supernatant was separated using Dynamag and subjected to immunoblotting.

#### Immunoblotting

Cells were lysed with ice cold RIPA lysis buffer (50mM Tris pH 8.0, 150mM NaCl, 1% Nonidet P-40, 0.5% sodium deoxycholate, 0.1% sodium dodecyl sulphate) with PMSF (Sigma, cat no. P7626) and proteinase inhibitor (Sigma, cat no. P8340). Cell lysates were separated on SDS-PAGE, blotted onto a Amersham Hybond P 0.45 poly-vinylidene fluoride (PVDF) membrane (GE healthcare, cat no. 10600023), and blocked with 5% bovine serum albumin (MP Biomedicals, cat no. 160069) or skimmed milk in 1× Tris-buffered saline (20mM Tris-HCl pH 7.4, 150mM NaCl) containing 0.1% Tween 20 (Sigma, cat no. P9416). Membrane was incubated in primary antibody (diluted in blocking solution) overnight at 4°C. This was followed by washing and incubation with the appropriate HRP-conjugated secondary antibody (goat anti-mouse HRP IgG H+L horse radish peroxide conjugate, Invitrogen, cat no.G21040, goat anti-rabbit IgG H+L horse radish peroxide conjugate, Invitrogen, cat no.G21234) for 1hour at room temperature. Protein bands were detected by applying either SuperSignal West Pico Chemiluminescent Substrate (Thermo Fisher Scientific, cat no. 34080) or SuperSignal West Femto Maximum Sensitivity Substrate (Thermo Fisher Scientific, cat no. 34095) to the membrane and imaging with Amersham Imager 600.

#### Immunostaining

Cells were cultured on glass coverslips coated with 0.2% gelatin and fixed with 4% paraformaldehyde (PFA) for 10 min. This was followed by blocking and permeabilization using 5% BSA containing 0.1% Triton X-100 (Sigma, cat no.T8787) and overnight incubation with primary antibody (diluted in blocking solution to the appropriate concentration) at 4°C. Next day, cells were washed with 1× PBS containing 0.1% Triton X-100 and incubated for 1 hr at room temperature with secondary antibody conjugated with fluorophore (Alexa fluor 488 donkey anti-mouse IgG H+L, Invitogen, cat no. A21202, Alexa fluor 555 goat anti-rabbit IgG H+L, Invitogen, cat no. A21428) diluted in blocking solution to appropriate concentration. Cells were washed, stained with DAPI, and mounted using VECTASHIELD (Vector Laboratories, cat no. H1000). All confocal imaging was done using the NIKON confocal microscope and imaged using a 60X 1.4 NA objective.

#### Super TOP-Flash assay

WT and *Cdh1^-/-^* mESCs were transfected with 400ng of TOP-Flash plasmid or 400ng of negative control FOP-Flash plasmid with Lipofectamine 2000. 25ng of pRL-TK (vector containing Renilla luciferase driven by HSV-thymidine kinase promoter) was also transfected together with the reporter plasmids for normalisation of the luciferase reporter. After transfection, the cells were treated with GSK3β inhibitor CHIR99021 (3µM, 6µM, 9µM). The assay was performed with the Dual-Glo Luciferse Assay System (Promega, cat no. E2940) according to the manufacturer’s protocol and luminescence was measured using Promega Glomax Luminometer. The firefly luciferase/ *Renilla* luciferase ratio was calculated for each sample.

#### Analysis of RNA-Seq data

Adapters were trimmed from FASTQ files using cutadapt version 2.10 (34, 35) with default settings keeping a minimum read length of 20 bp. Reads were mapped to the GRCm38 mouse genome keeping only uniquely mapping reads using STAR version 2.7.5b (36) with the following settings “--outFilterMultimapNmax 1”. Reads overlapping genes were then counted using featureCounts version 2.0.1 (37, 38) using the Gencode version 25 basic transcriptome annotation. The count matrix was imported into R. Only genes with at least 1 count per million (CPM) across at least 3 samples were kept. Differentially expressed genes between *Cdh1^-/-^* and WT control samples were then identified using limma version 3.44.3 (39). Significant differentially expressed genes were defined as having an FDR adjusted p-value < 0.05. The raw sequencing files generated during this study are available at GEO: GSE180562.

#### qRT PCR

Total RNA was isolated using TRIzol (Life Technologies, cat no. 15596018) and was quantified using NanoDrop Spectrophotometer followed by Dnase treatment. cDNA was made from 1µg of RNA using Verso cDNA Synthesis Kit (Thermo Scientific, cat no. AB-1453/B). cDNA was then diluted 1:10 and 1µl of it was used for qRT-PCR along with gene-specific primers (Supp. Table 4) and Power SYBR Green PCR Master Mix (Applied Biosystems, cat no. A25742). *Gapdh* was used as endogenous control. Data were analysed after normalising the expression of the particular gene to that of *Gapdh*.

#### Embryoid body formation

Embryoid bodies were formed by hanging drop method. 500 cells were suspended in a 20µl drop of ESC media (15% FBS) without LIF and incubated at 37°C and 5% CO_2_ for 72hours. Post 72hours, these embryoid bodies were cultured on 0.2% gelatin coated tissue culture dishes in DMEM medium containing 10% FBS for 5 days, followed by collection for protein lysates or isolating RNA.

#### Fitting linear mixed-effects models

We used linear mixed-effects models (Matlab function fitlme) to examine the influence of 3 factors (*Cdh1* status, cell differentiation status and GSK3β inhibitor status) on the expression levels of β-CATENIN targets. Each of the 3 factors, all possible 2-way interactions between these 3 factors and a 3-way interaction between the factors, were included as predictors for the model. Repeat number was included as a random effect. Adjusted r^2^ values were used as indicators of the quality of the fit (Supp. Table 1). Predictors were considered significant at a significance level of 0.05. Residuals obtained for the models were visually examined to ensure homogeneity and normality. In addition, normality of residuals was tested using the Anderson Darling test (Matlab function adtest) and was found to be non-significant for 12/13 genes (p > 0.05).

#### Principal component analysis

To examine the expression of all of the 13 genes in different experimental conditions, we considered each gene as one variable and carried out principal component analysis. For each gene, expression levels relative to *Gapdh* were natural log-transformed and converted into z-scores by subtracting the mean of all experiments and dividing by the standard deviation across all experiments. Each experimental condition was represented by a row vector that included the normalized expression of each of the 13 genes. Principal components analysis was carried out using the Matlab function pca. The first 6 principal components (PCs) explained 94.5% of the variance. The first 2 PCs explained 68% of the variance and were used to visualize gene expression differences across experiments.

## Supplementary Figure Legends

**Supplementary Figure 1:**
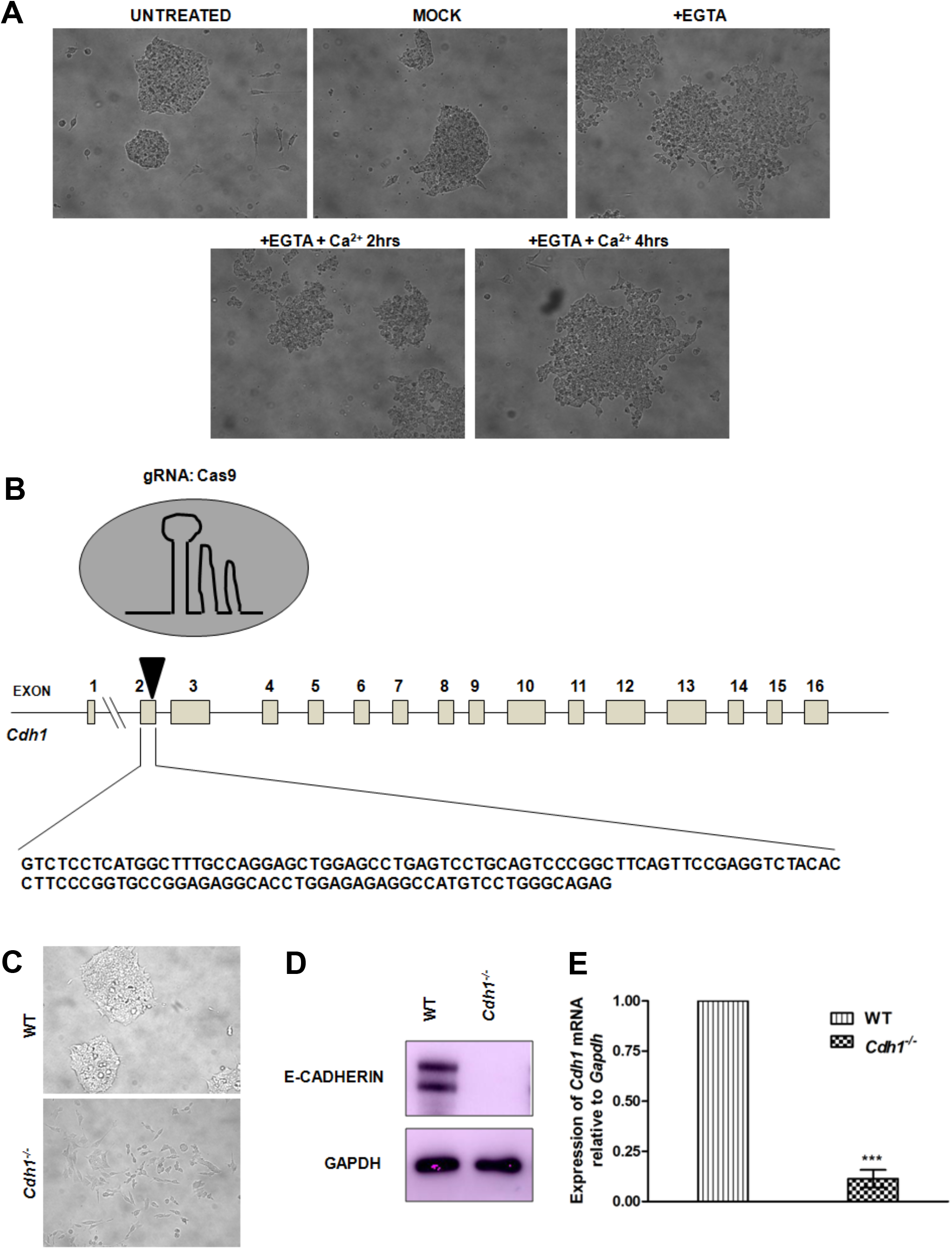
A. Bright field images of WT mESCs after treatment with EGTA, and subsequent restoration with Ca^++^ for the indicated time. Untreated and mock-treated samples were used as controls. B. The strategy for knocking out *Cdh1* using CRISPR-Cas9 technology is shown with the target site of the sgRNA indicated by a black triangle. C. Bright field images showing morphology of WT and *Cdh1^-/-^* mESCs. D. Western blot showing expression of E-CADHERIN in *Cdh1^-/-^* mESCs compared to WT. E. Graph showing relative expression of *Cdh1* in *Cdh1^-/-^* mESCs compared to WT. Error bars represent mean ± SD from three independent experiments. *p<0.05; **p<0.01; ***p<0.001 by two-tailed Student’s t-test.

**Supplementary Figure 2:**
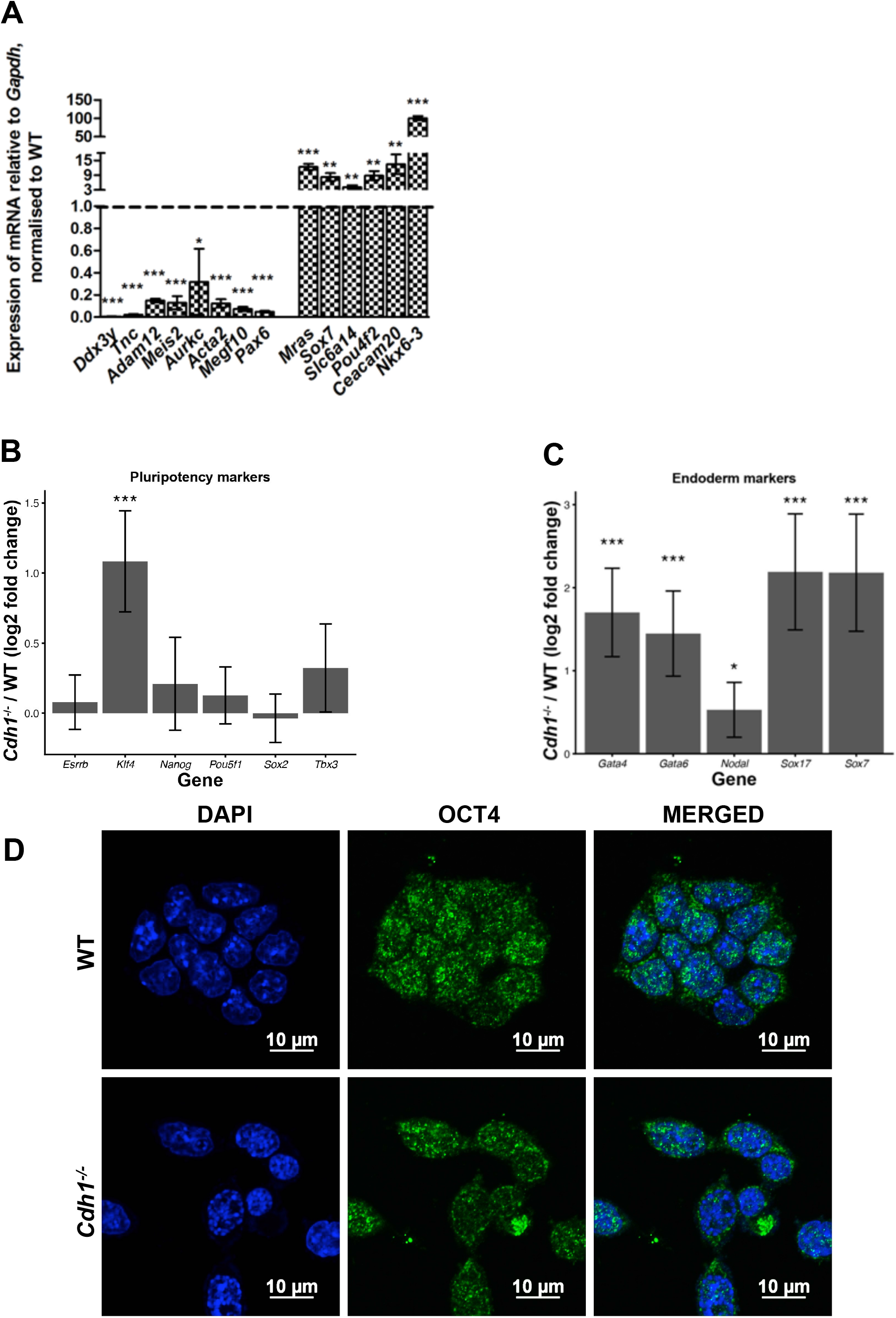
A. Graph representing the expression of the highly downregulated and the highly upregulated genes in WT and *Cdh1^-/-^* mESCs as validation of the RNA-seq data. Gene expression is relative to *Gapdh* and relative to WT (n=3). B, C. Graphical representation depicting the fold change in the expression of (B) pluripotency factors, and (C) endoderm markers at the transcript level in *Cdh1^-/-^* mESCs compared to WT by RNAseq analysis. D. Representative confocal image showing expression of OCT3/4 in WT and *Cdh1^-/-^* mESCs. Scale bar, 10µm. For all experiments, error bars represent mean ± SD from three independent experiments. *p<0.05; **p<0.01; ***p<0.001 by two-tailed Student’s t-test.

**Supplementary Figure 3:**
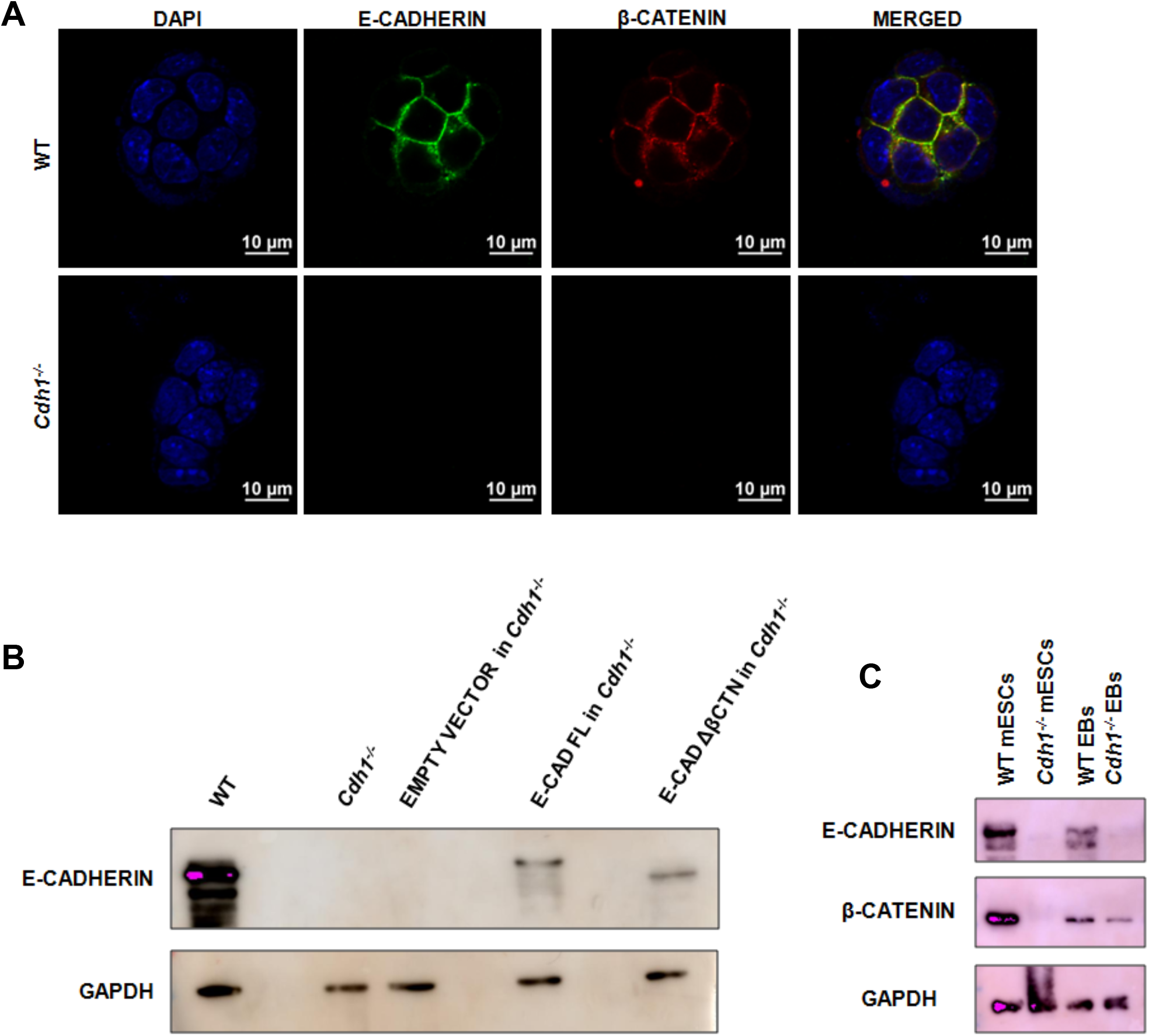
A. Representative confocal images showing expression of E-CADHERIN and β-CATENIN in WT and *Cdh1^-/-^* mESCs. Scale bar, 10µm. B. Western blot confirming the exogenous expression of E-CADHERIN in *Cdh1^-/-^* mESCs stably transfected with E-CAD FL, E-CAD ΔβCTN. C. Western blot showing levels of E-CADHERIN and β-CATENIN in WT and *Cdh1^-/-^* mESCs and embryoid bodies (EBs).

**Supplementary Figure 4:**
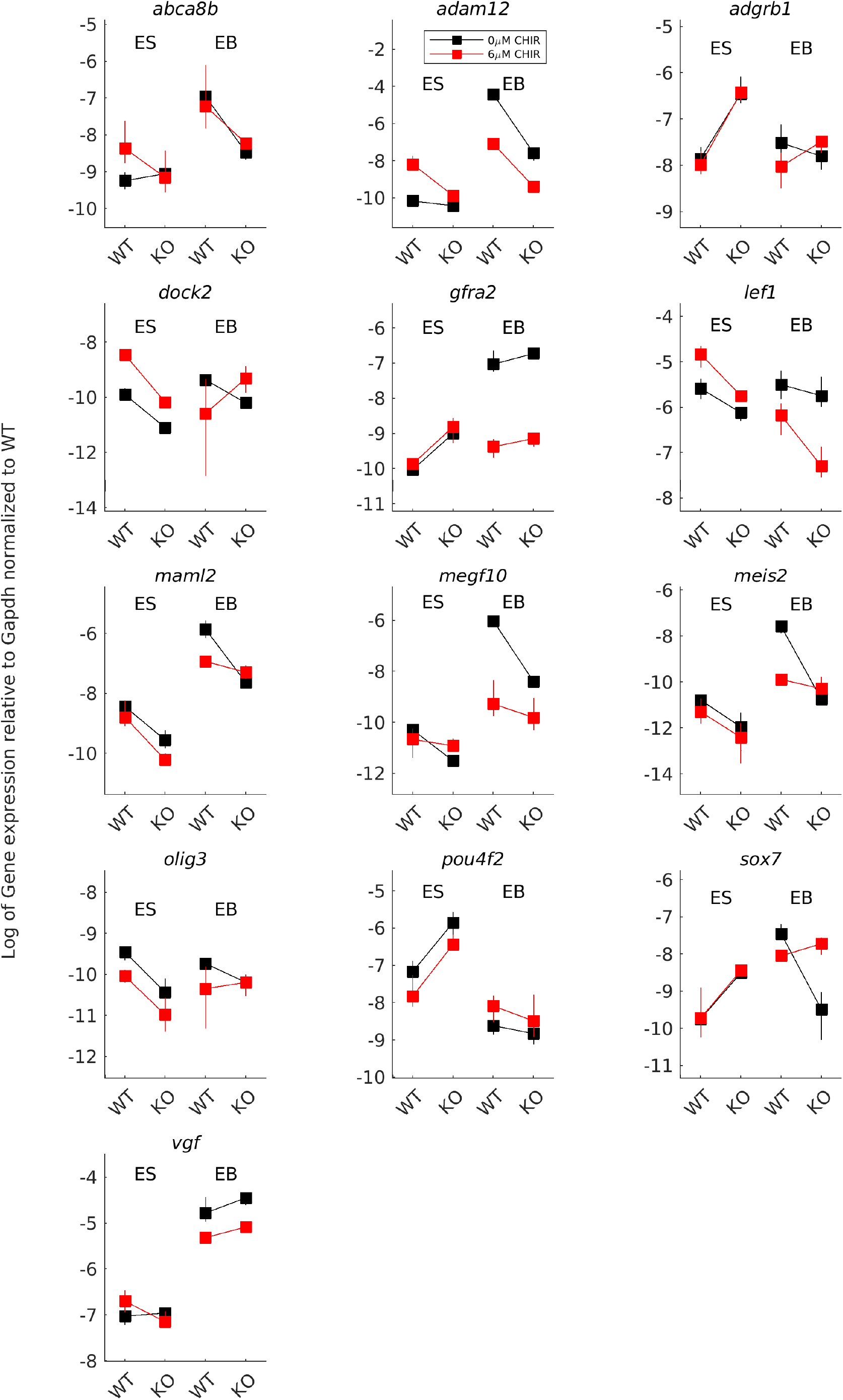
Each plot shows the expression of one of the 13 targets of β-CATENIN in WT and *Cdh1^-/-^* mESCs and EBs. Black lines represent cells treated with DMSO and red lines represent cells treated with 6µM CHIR99021. Squares and whiskers represent means and 95% confidence intervals across 3 repeats. The natural logarithm of expression levels shown are relative to *Gapdh* and normalized to WT in that plate.

**Supp. Table 1:**
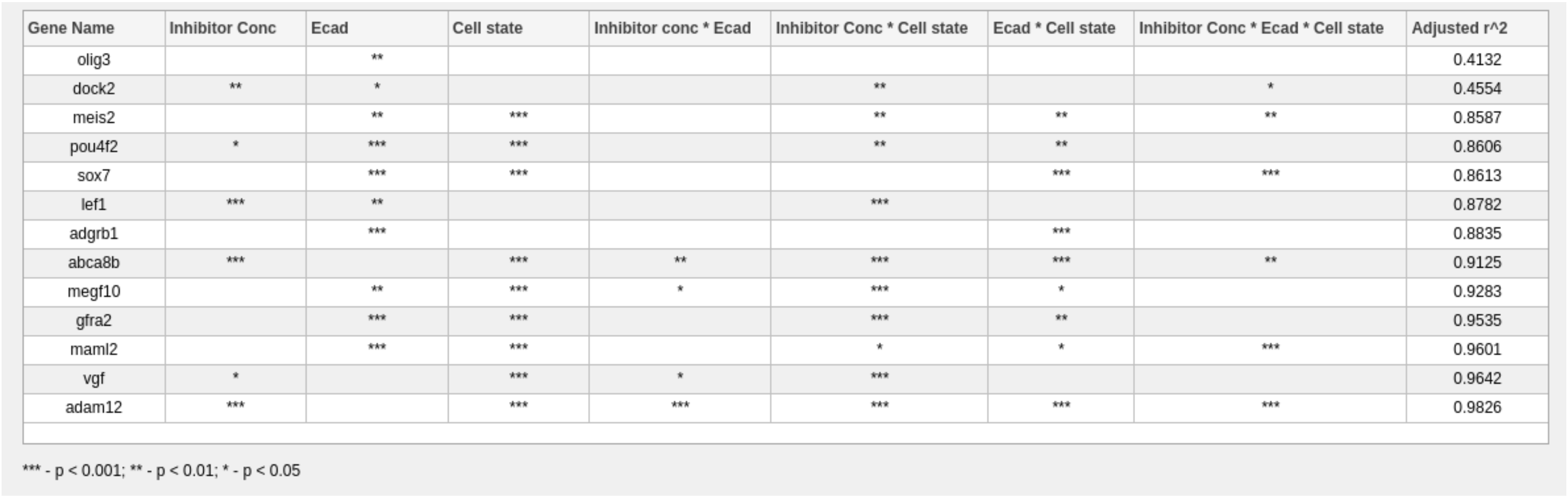
Table shows the significant predictors for each gene identified using a linear mixed-effects model. The columns represent the different predictors used as part of the model. Predictors included individual predictors as main effects, all possible 2-way interactions and one 3-way interaction. The last column show the adjusted r2, an indicator of the quality of model fit.

**Supp. Table 2: Table showing.** the co-efficients for the first 6 principal components (these 6 accounted for 94.51% of the total variance). Significant co-efficients are shown in bold (significance was calculated as values that are greater than the value expected if all 13 genes contributed equally, i.e if co-efficient was < -1 / square root of (13) or > 1 / square root of (13).

| Gene Name | PC1 | PC2 | PC3 | PC4 | PC5 | PC6 |
| --- | --- | --- | --- | --- | --- | --- |
| abca8b | **0.3225** | 0.0727 | -0.0086 | **0.3511** | -0.2061 | 0.0993 |
| adam12 | **0.3551** | -0.0135 | **0.2834** | 0.0706 | -0.2016 | 0.0602 |
| adgrb1 | -0.2046 | **0.4206** | **0.3059** | 0.1794 | **0.4019** | 0.0296 |
| dock2 | 0.1202 | **-0.4248** | -0.0420 | 0.2739 | **0.5009** | **0.6495** |
| gfra2 | 0.2342 | 0.2074 | **0.4785** | **-0.3430** | **0.4016** | -0.1016 |
| lef1 | -0.0076 | **-0.4323** | **0.6381** | 0.1339 | **-0.2866** | -0.0205 |
| maml2 | **0.3718** | 0.0215 | -0.1993 | 0.0331 | -0.0901 | -0.1680 |
| megf10 | **0.3661** | 0.0329 | 0.2198 | -0.0056 | 0.1131 | -0.0847 |
| meis2 | **0.3503** | 0.0311 | 0.0043 | 0.2132 | -0.1224 | -0.1304 |
| olig3 | 0.1503 | **-0.4155** | -0.1977 | 0.1768 | **0.4562** | **-0.6284** |
| pou4f2 | **-0.3143** | 0.0921 | 0.1897 | **0.4265** | 0.0485 | **-0.3080** |
| sox7 | 0.1656 | **0.4460** | -0.1141 | **0.5053** | 0.0403 | 0.0833 |
| vgf | **0.3389** | 0.1570 | -0.1094 | **-0.3375** | 0.1144 | 0.0626 |

**Supp. Table 3:**
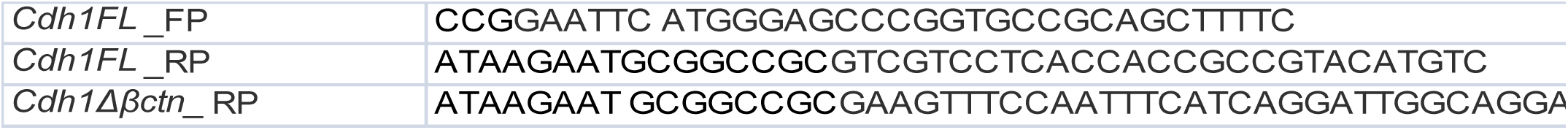
List of primers used for cloning

**Supp. Table 4:**
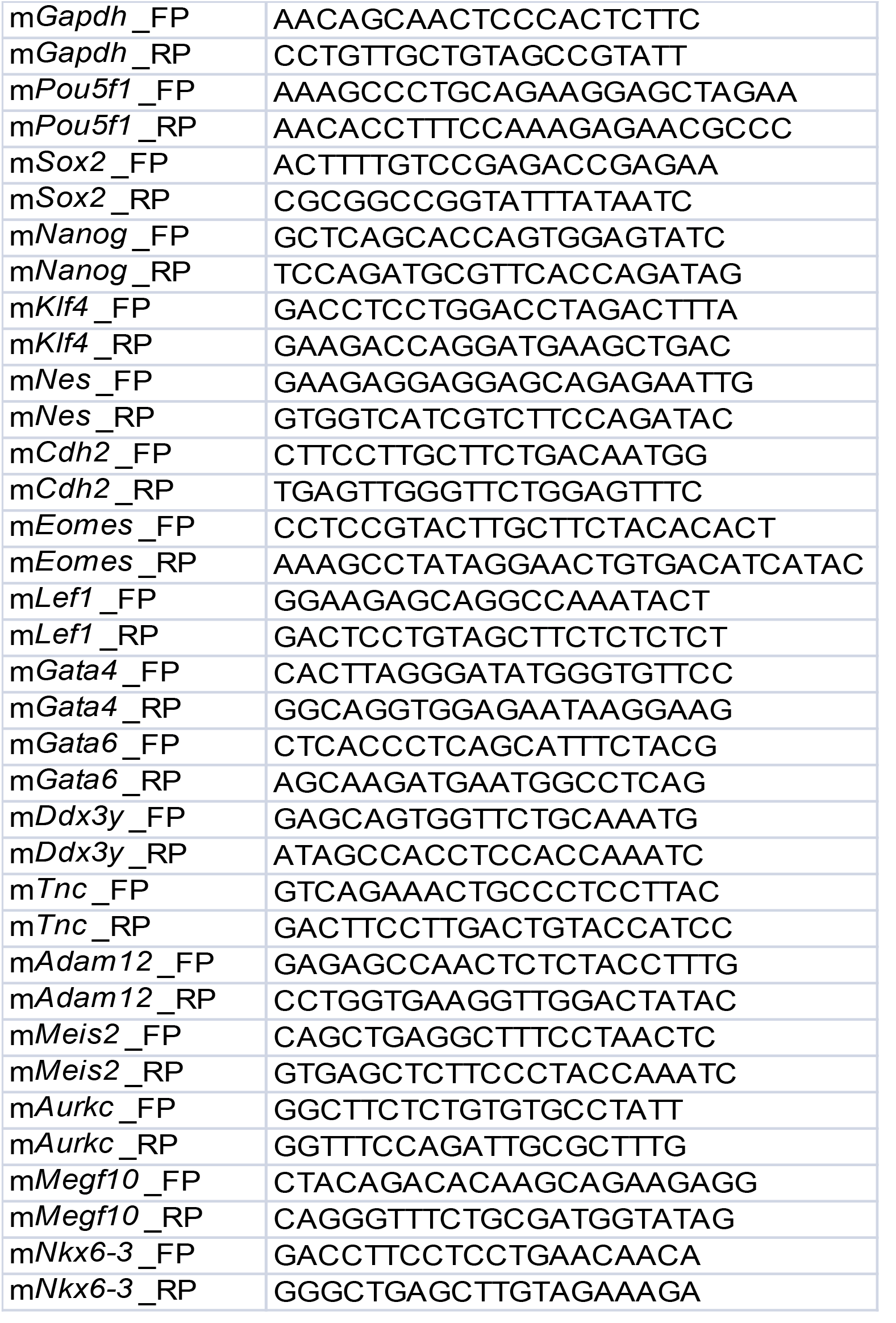
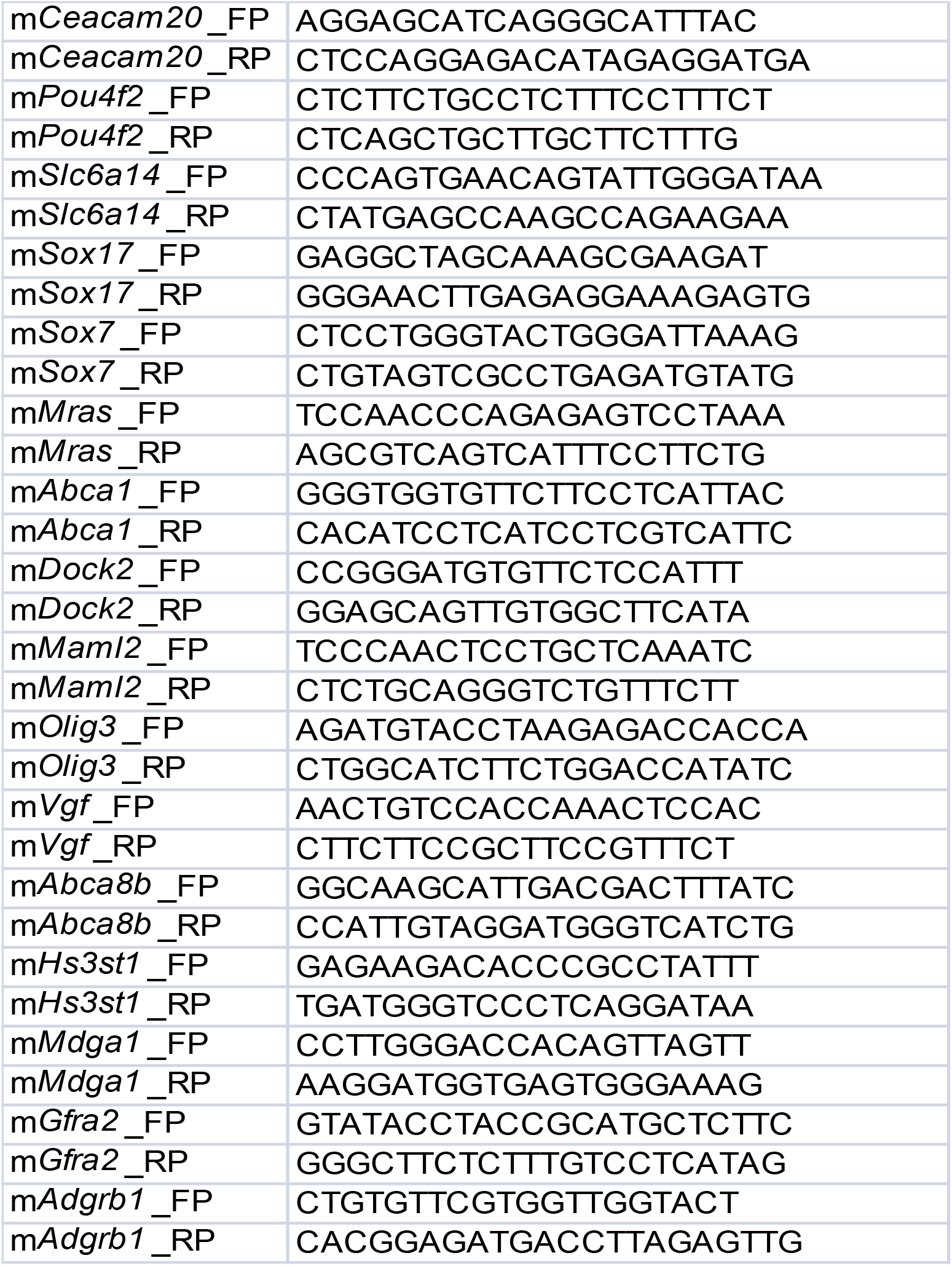
List of primers used for qRT PCR:

